# Influence of mixed and single infection of grapevine leafroll-associated virus and viral load on berry quality

**DOI:** 10.1101/2023.11.15.567278

**Authors:** Wisam Salo, John A. Considine, Michael J. Considine

## Abstract

Grapevine leafroll disease (GLD) is a viral disease that affects grapevines (*Vitis vinifera* L.) and has a severe economic impact on viticulture. In this study, the effect of grapevine leafroll-associated viruses (GLRaV) on berry quality was investigated in clones of cultivar cv. Crimson Seedless table grapes infected with GLRaV. RT-PCR confirmed the identity of the clones: clone 3236, infected only with GLRaV-3 (termed Single); clone 3215, infected with GLRaV-3, GLRaV-4 strain 9 and grapevine virus A (termed Mixed), and a viral free clone of the same genetic background of the infected clones (termed Control). The berry quality indices of size, sugar, acidity, and anthocyanin content were measured at harvest maturity. RT-qPCR was used to determine viral load. The study was repeated over two years. A two-way, multivariate analysis of variance (MANOVA) was applied with clone and season as independent variables and the measured berry quality parameters as a dependent variable. All dependent variables were significantly affected by viral infection (Wilks, λ, [2,33] = 0.033895, p-value < 0.001), while only titratable acidity (TA) was affected by season. Average berry dry mass decreased (p-value < 0.001). The water content of both infected clones was greater than that of the control (p-value < 0.001). Both infected clones displayed reduced sugar content as a fraction of the berry dry mass (p-value < 0.001). The anthocyanin and the phenol content of the infected clones were significantly reduced compared to the control clone (*p* < 0.001, *p* < 0.05, clone 3236 and clone 3215, respectively). Finally, the viral load was highly variable, and no quantitative relationship between viral load and berry composition was found.

## 2 Introduction

It is well known that the cultivated grapevine (*Vitis vinifera* L.) is exceptionally vulnerable to viral infection. Grapevine leafroll-associated viruses (GLRaV) are among the most widespread in vineyards (Martinson et al. 2008; Naidu et al. 2014). It has been established that six species of the family Closteroviridae are responsible for grape leafroll disease (GLD) (Adiputra et al. 2019; Maree et al. 2013; Sharma et al. 2015; Velasco et al. 2014). This disease may result in a severe reduction in fruit yield, vigour and a delay in fruit ripening of wine grapes (Cabaleiro et al. 1999; Naidu et al. 2014). Although extensive studies have been conducted, the mechanism of virus-host interaction that affects the impact on fruit quality is still uncertain. In addition, the majority of reports were conducted with red-berried wine varieties (Alabi et al. 2016; El Aou-ouad et al. 2016; Lee et al. 2009; Lee and Martin 2009; Montero et al. 2016a). Comparatively, few studies have been conducted on table grapes, where the effects on quality and yield are less pronounced or deleterious (Singh Brar et al. 2008). To date, studies of GLD have focused on a single virus species, predominantly GLRaV-3, because of its high impact and virulence. In comparison, the influence of GLRaV-4 strain 9 on berry quality in grapevine has scarcely been reported (Maree et al. 2013), nor has the impact of mixed infections. In both wine and table grapes, viral infection appears to delay ripening (Komar et al. 2010; Lee et al. 2009; Lee and Martin 2009), leading to reduced sugars and anthocyanins and an increase in berry weight and volume. Alabi et al. (2016) indicate that the viral effect on sugar levels was more significant at and after véraison than pre-véraison. Thus, changes in berry morphology may be linked to source-sink effects since GLRaV-3 spreads through the phloem and post véraison increase is due almost wholly to the mass flow of phloem solution (Choat et al. 2009; Wang et al. 2003; Zhang and Keller 2017). This study sought to investigate the nature of the influence of single (clone 3236) and mixed (clone 3215) viral infections on the berry quality of cv. Crimson Seedless *Vitis vinifera* L. as an example of table grapes grown under commercial vineyard conditions. Further, it sought to determine whether the viral copy number influenced berry quality.

## 3 Material and Method

### 3.1 Plant material

The study utilised two clones of *Vitis vinifera* L. cv. Crimson Seedless was previously generated by the Department of Agriculture and Food of Western Australia and was used in a previous study. Clone 3236, infected with GLRaV-3 with mild symptoms, will represent the single infection group. Clone 3215, infected with GLRaV-3, GVA, and GLRaV-4 strain 9, will represent the mixed infection group (Alagappan 2011; Singh Brar et al. 2008). Both infected clones were compared to cv. Crimson Seedless viral-free vines (mock-infected). Each group had six vines as a biological replicate. The vines were 15 years old, grafted onto Schwarzmann rootstock and grown in a commercial vineyard located in the Swan Valley in Western Australia (-31.827789, 115.999947). The grapevines were spaced 3.3 m between the rows and 2.4 m between the vines. The infected vines were set in one row in the vineyard; each clone was planted in three replicates, followed by three vines of the other clone, and separated by a healthy vine. The control vines were located in the left adjacent row of the infected clones. The control clones have been checked during two seasons of the study for the presence of the disease symptoms and RT-PCR test has been carried out to confirm the absence of viral infection in the vines. Then the result was confirmed with qPCR in the experiment of the viral load. Moreover, no infection has been reported with other types of viruses to the Department of Primary Industries and Regional Development of WA during the two seasons of the study.

### 3.2 Berry Sample Collection

Five bunches per vine were randomly collected at the harvest stage of two seasons (EL38, March 2017/18). All the samples were collected on the same day and finished before 10:00 am. Finally, °*Brix* and titratable acidity were measured on the sampling day. The ripening stage of the berries was confirmed using the recommendation of OIV resolution VITI 1/2008 (OIV 2008) and UE Commission Regulation 543/2011. Where table grapes are considered to be ripe at °*Brix* value higher than 16 ^⍰^*Brix* or when the SSC (expressed as g.L^-1^)/TA (expressed as g.L^-1^ tartaric acid) ratio is higher than 20; with regards to the exception the case of seedless varieties, ripeness is considered at TSS 14 ^⍰^*Brix*. All bunches were checked to be free from any fungus symptoms and any other physical damage. The bunches of each vine were loaded into plastic bags and immediately kept on ice. Randomly, 50 berries were selected per vine, weighed and then macerated for 5 minutes in a blender. As demonstrated by Peppi et al. (2006), the filtered juice was used for soluble solids and titratable acidity measurements. An extra ten berries were kept for dry mass measurement.

### 3.3 Viral status

Virus identification was carried out by reverse transcription PCR using previously designed primers, were the GLRaV’s characterised using hHSP70 (Osman and Rowhani, 2006, Osman et al. 2007). The GVA virus was characterised by target the cap protein (Minafra and Hadid 1994). The 18srRNA gene used as internal control (Minafra and Hadidi 1994). Five petioles, free from any symptoms of fungus, mould, and other physical damage, were randomly sampled per vine at the berry pea-size stage (EL31) (Coombe and Iland 2004) December 2016. RNA was extracted using the Spectrum™ Plant Total RNA Kit (Sigma-Aldrich, St. Louis, MO, USA) according to the manufacturer’s instructions. DNA was removed by applying the ON-COLUMN DNASE I DIGESTION SET (Sigma-Aldrich, St. Louis, MO, USA). The average concentration for the extracted RNA was 92-150 μg.μL^-1^. The RNA was diluted to 2-5 μg.μL^-1^, and cDNA synthesis was performed using the SuperScript™ IV VILO™ Master Mix (Invitrogen, Carlsbad, CA, USA) following the manufacturer’s instructions. Previously designed primers were selected for detecting the viruses. For GLRaV-3, the pairs Lc1/F and Lc2/R have been used (Osman and Rowhani 2006). For GLRaV-4 strain nine and five, the pairs LR9/F, R and LR5HSPC/F, R respectively (Osman et al. 2007). Those sets were targeting the hHSP70 gene in the leaf roll viruses. On the other hand, the CP gene was targeted in the detection of GVA virus using the primer pair GVAC1/F, R that was predicted by Minafra and Hadidi (1994). In addition, 18S rRNA was used as a positive control (Gambino and Gribaudo 2006). The reaction mixture and the reaction parameters were modified after optimisation as 5 μL GoTaq Green Mastermix (Promega, Madison, USA) 0.5 μL of 10 mM of each primer, 1.5 μL nuclease-free water and 2.5 μL of the cDNA. The amplification steps were: 2 min at 94 °C, 35 cycles of 94 °C for 30 s, 58 °C for 45 s, and 72 ⍰C for 60 s, and a final extension at 72 °C for 7 min. The electrophoresis was carried out by loading 10 μL of the amplification mix in a 1.5% agarose gel submerged in TAE buffer (40 mM Tris base, 20 mM sodium acetate, 1 mM EDTA pH 8.0). The amplified DNA fragments were visualised on a UV transilluminator following ethidium bromide staining and photographed. Positive and negative controls for the viruses under study were included in each experiment (Sambrook and Russell 2001).

### 3.4 Viral Load

The rachis with pedicels attached (here-after, “stalk”) were collected with the berries at the harvest stage (EL38, March 2017/18). The RNA was extracted as described for viral identity, with the exception that traces of DNA were removed using Dnase I Amplification Grade (Sigma-Aldrich, St. Louis, MO, USA) according to the manufacturer’s instructions. The cDNA synthesis was carried out as described in the manufacturer’s instructions of SuperScript™ IV VILO™ Master Mix (Invitrogen, Carlsbad, CA, USA). Then, the cDNA was diluted to supply 10 ng in a 10 μl reaction volume to be subjected to the ABI 7500 Realtime PCR System (Applied Biosystems, Foster City, CA, USA) using PowerUp™ SYBR™ Green Master Mix and previously designed primers (Osman and Rowhani 2006; Osman et al. 2007, 2008). 18S rRNA used as a housekeeping gene (Osman et al. 2008). The reaction was prepared according to the manufacturer’s instructions.

The positive controls were constructed as plasmid vectors and contained the specific region of hHSP70 for each of the leafroll viruses, CP for the GVA virus and 18S rRNA as a housekeeping gene. These reigns are short sequences between 240 and 280 bp that contain the primer sites. They were chosen from the virus reference genomes on the National Centre for Biotechnology Information website (https://www.ncbi.nlm.nih.gov/). The accession numbers are KY821094.1 (GLRaV-3), AY297819.1 (GLRaV-4/9), AF039552.1 (GLRaV-4/5), AY244516.1 (GVA) and AF321271.1 (18s rRNA). The designed vectors were aligned and synthesised by Integrated DNA Technology Australia (IDT) (NSW, Australia), providing the PUC IDT-AMP vectors and the vector construction process. All the vectors were supplied in 4 μg lyophilised powder, and the stock solution was prepared by adding 40μl of nuclease-free water directly to the lyophilised powder to get 100 ng.μL^-1^. The working solution with a specific concentration (10 ng.μL^-1^) was prepared by taking 3 μL from the stock solution (100 ng.μL^-1^) and diluting it in 27 μL nuclease-free water. The copy number per 1 ng was determined by using the equation (1).

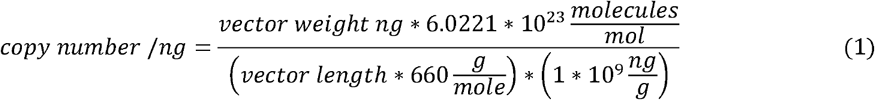

The standard curves were prepared for all viruses using the working solution to produce ten-fold serial dilutions for 5-6 points in triplicates (102 to 107 copies). The concentration of each dilution was measured by Qubit 4 Fluorometric Quantification and RNA Quantification, broad range Kit (Thermo Fisher Scientific Australia Pty Ltd, VIC, Australia) and then detected by RT-qPCR in two independent assays per virus. Standard curves for each virus were constructed by plotting Ct values versus the logarithm of the RNA copy number (Ct vs the log of the standard sample amount) using the StepOne Software (Applied Biosystems, Foster City, CA, USA).

### 3.5 Berry quality

Total soluble solids (TSS): The juice filtered through the muslin cloth into the conical flask to exclude berry flesh debris. Finally, TSS was determined using a digital refractometer calibrated in °*Brix* (g sucrose/100 g solution, Atago, PAL-1 Digital *Brix* Refractometer, Tokyo, Japan). Titratable acidity (TA): The juice (10 ml) was titrated against 0.1 M NaOH to an endpoint pH8.2. TA was calculated using the formula and expressed as a percentage of tartaric acid equation (2) (Considine and Frankish,2013; Iland 2004).

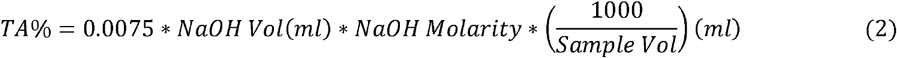

°**Brix/acid ratio**: For each sample, the °*Brix* to acid ratio was calculated by dividing the °*Brix* value by the percentage acidity. **Berry volume**: The berry’s size was determined. Callipers were used to determine the vertical diameter (L) and horizontal diameter (l). The volume was determined by matching the berry form to an ellipsoid using the following equation: volume (cm^3^) = 4 abc/3, where a = b = l/2 and c = L/2 (Río Segade et al. 2013; Río Segade et al. 2011). **Dry weight and water content**: All treatments are arranged into a set of three replicates for each sample. Each replicate contains 10 berries placed in a Petri dish. The initial weight of the samples and the Petri dishes were recorded. Then, the dishes were subjected to 57^°^C for 21 days in a dry vacuum oven. The weight of all treatments was measured every day until no change in weight was noticed. The dry weight was calculated as illustrated in equation (3), while the total moisture in berries was calculated according to equation (4) (Nielsen 2017):

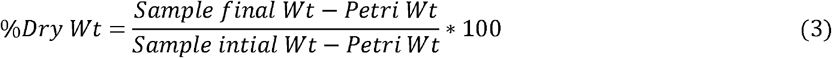

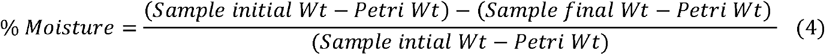

### 3.6 Total anthocyanin and phenol determination

The frozen berries were thawed at 4^°^C for 2hr. The accurate weight of 50 berries per sample was recorded. The berries were transferred to a 50 mL plastic beaker for homogenisation using an Ultra-Turrax® T25 high-speed homogeniser with an S25N dispersing head (Janke and Kunkel GmbH Co. Germany). The beaker was placed on ice and homogenised at 24,000 rpm for 30 sec. The shaft was cleaned, and the remaining grape tissue was returned to the homogenisation vessel again and homogenised for 15 s. One g of the homogenate was placed in a 15 mL tube. 10 ml of 50 % (v/v) aqueous ethanol pH2 was added. The tube was mixed by inversion for 1 hr and centrifuged at 3,500 rpm for 5 min. The supernatant was collected, and the volume was measured. One ml of the extract was taken and mixed with 100 ml of 1 M HCl and mixed thoroughly. The diluted extract was incubated at room temperature for 3 hr. The absorbance of the acidified diluted extract was measured at 520 nm using a 1.0 M HCl blank on a SPECTROstar Omega reader (BMG LABTECH, Ortenberg, Germany). The anthocyanin mg/berry was calculated (equation (5)) along with total phenolics per berry (absorbance units (au) per berry) as predicted in equation (6) (Iland 2004).

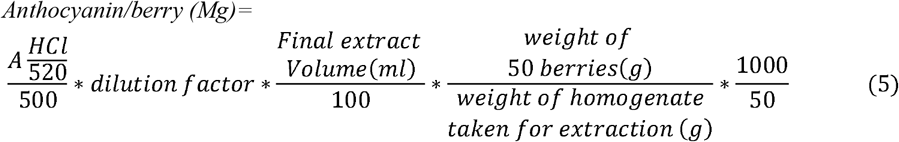

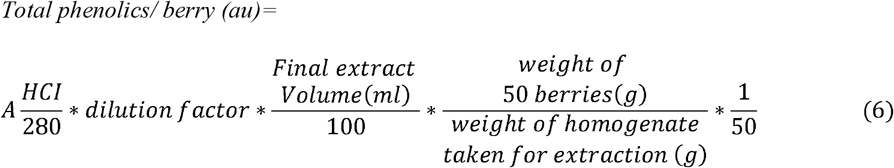

### 3.7 Statistical Analysis

The statistical analysis was performed using the “R” language (https://cran.r-project.org/). A two-way multivariate analysis of variance (MANOVA) was conducted using the viral infection type and seasonal influence as independent factors and the measured berry quality characteristics as dependent variables. Tukey post-hoc contrasts have been used to analyse the difference between the means of berry quality parameters and the clones. Furthermore, a simple linear regression was calculated to predict the relationship between the berry mass and sugar mass, titratable acidity (TA), and total soluble solids (TSS). Finally, the ANCOVA test was applied to analyse the viral load and seasons against the berry quality parameters. The figures have been plotted using the R package “ggpubr” (https://CRAN.R-project.org/package=ggpubr)

## Results

### Viral identity

All the samples showed a positive PCR product of 844bp, representing the internal control 18S rRNA (Figure 1). The Control vines failed to amplify PCR products using gene-specific primers for GLRaV-3, GLRaV-4 strain 9 and GVA, confirming the absence of detectable infection of these viruses. The Clone 3236 vines tested positive to GLRaV-3, but not GLRaV-4 strain 9 or GVA. The clone 3215 vines consistently tested positive for GLRaV-3 and GVA, which confirmed the mixed infection. However, only four vines amplified bands with the GLRaV-4 strain 9-specific primers. Vines that do not show amplification to GLRaV-4 are used as a mixed; hence they still have an infection with GLRaV-3 and GVA. GVA has been mostly related with GLRaV-1 and -3, with the hypothesis that co-infection contributes to the severity of grape symptoms. Previous research (Credi and Babini 1997) predicted that disease severity increased with mixed infections in grapevine, particularly when GVA co-infected with GLRaV-3.

**Figure 1.**
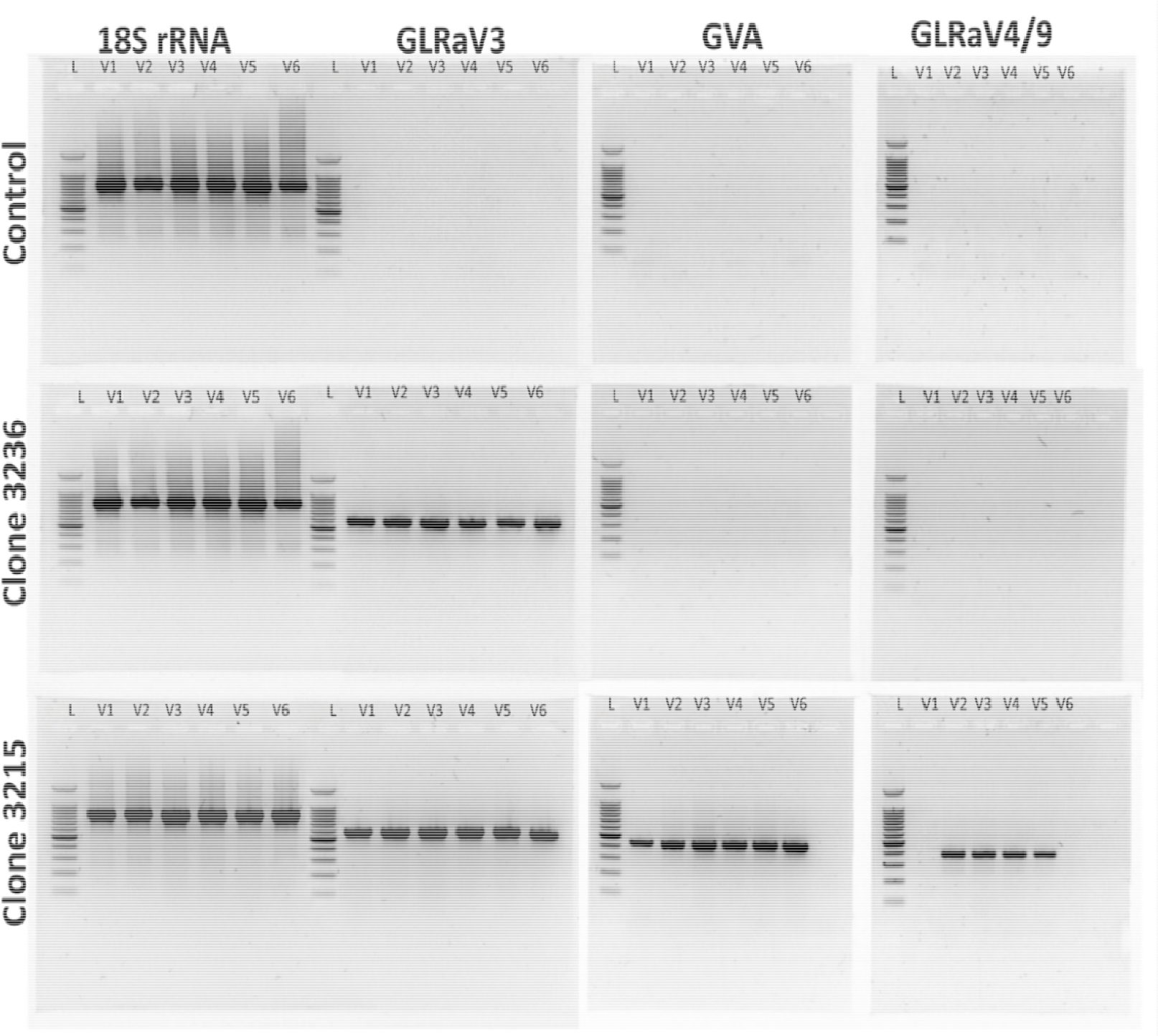
Agarose gel electrophoresis RT-PCR amplified cDNA products of GLRaV-3 GLRaV-4 Strain 9, GVA and 18s rRNA. L: Ladder DNA Promega 100bp with fragment size 100 to 12000 bp. V1-V6 represent the vines replicates of each clone. Vertically: 18S rRNA represented the amplification of the positive control. GLRaV-3 represent the amplification of HSP70 of the GLRaV-3 (546bp). GVA represent the amplification of CP of the GVA (429bp) GLRaV-4 strain 9 represent the amplification HSP70 of GLRaV-4 strain 9 (393bp), in the Control, Clone 3236, and Clone 3215.

### Berry quality

Fruits on infected vines were lighter in colour and larger than those growing on uninfected vines (Figure 2). Having established the identity of viral infections in the vines, berry quality parameters were quantified for two seasons (2017/18). A two-way multivariate analysis of variance (MANOVA) was performed with the viral infection type and the seasonal effect as the independent variables and the quantified berry quality parameters as dependent variables. The viral infection types included three levels viral free (Control), single infection (GLRaV-3) and mixed infection (GLRaV-3, GLRaV-4 strain 9 and GVA), while the seasonal effect consisted of two levels (2017 and 2018). Findings show significant differences for all dependent variables (Wilks λ, [2,33] = 0.033895, p < 0.001) for the viral infection type, while no significant difference was observed for the seasonal effect, with the exception of TA (Table 1).

**Table 1.**
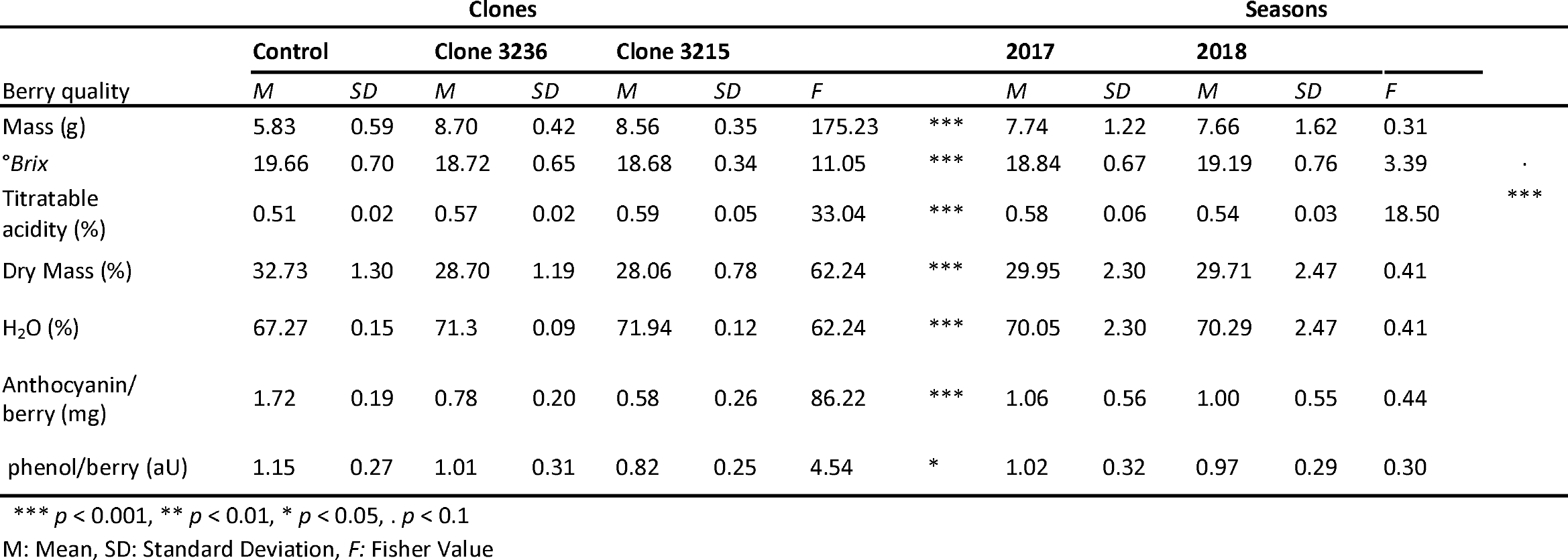
The variation of the berry quality of the infected clones compares with the control. Means, standard deviations, Fisher value and the *p* Value were calculated using MANOVA, where each clone comprise 6 replicates (n=6).

**Figure 2.**
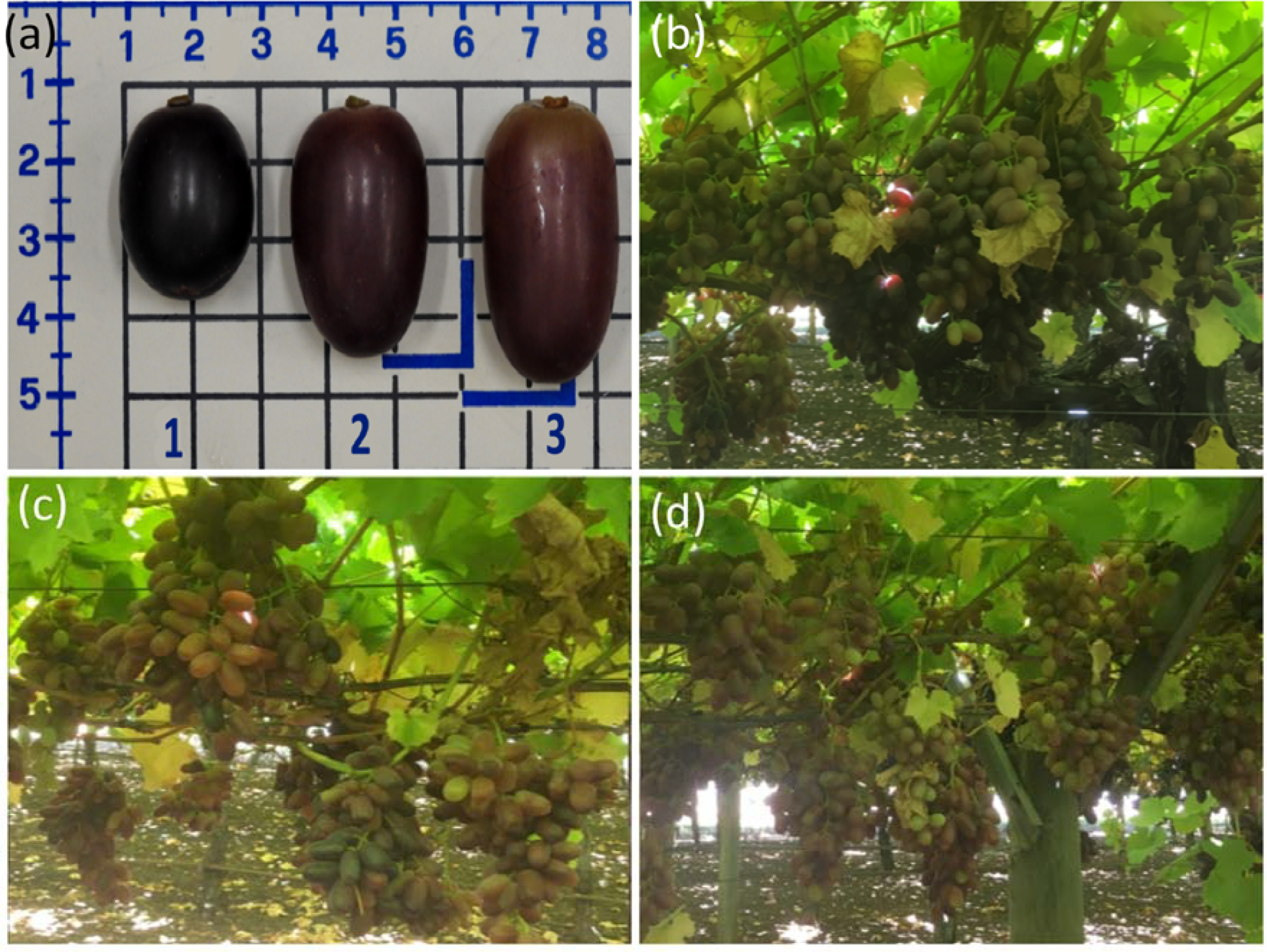
The cv. Crimson Seedless berries on the harvest day, showing the three types of viral infection: **(a)** shows the individual berries at the harvest stage, where number 1 represents the control, number 2 represents clone 3236 and number 3 represents clone 3215. Figure **(b), (c)** and **(d)** shows the corresponding berries on the control vine, clone 3236 and clone 3215 respectively, immediately prior to harvest.

The mean of the berry mass was strongly dependent on the presence of the infection but was unaffected by season. Tukey post-hoc contrasts showed significant differences between the Control and Single infection groups (*p* <0.000001) and the control and the Mixed infection (*p* <0.0001), but not between the two infection groups (Figure 3(a)).

**Figure 3.**
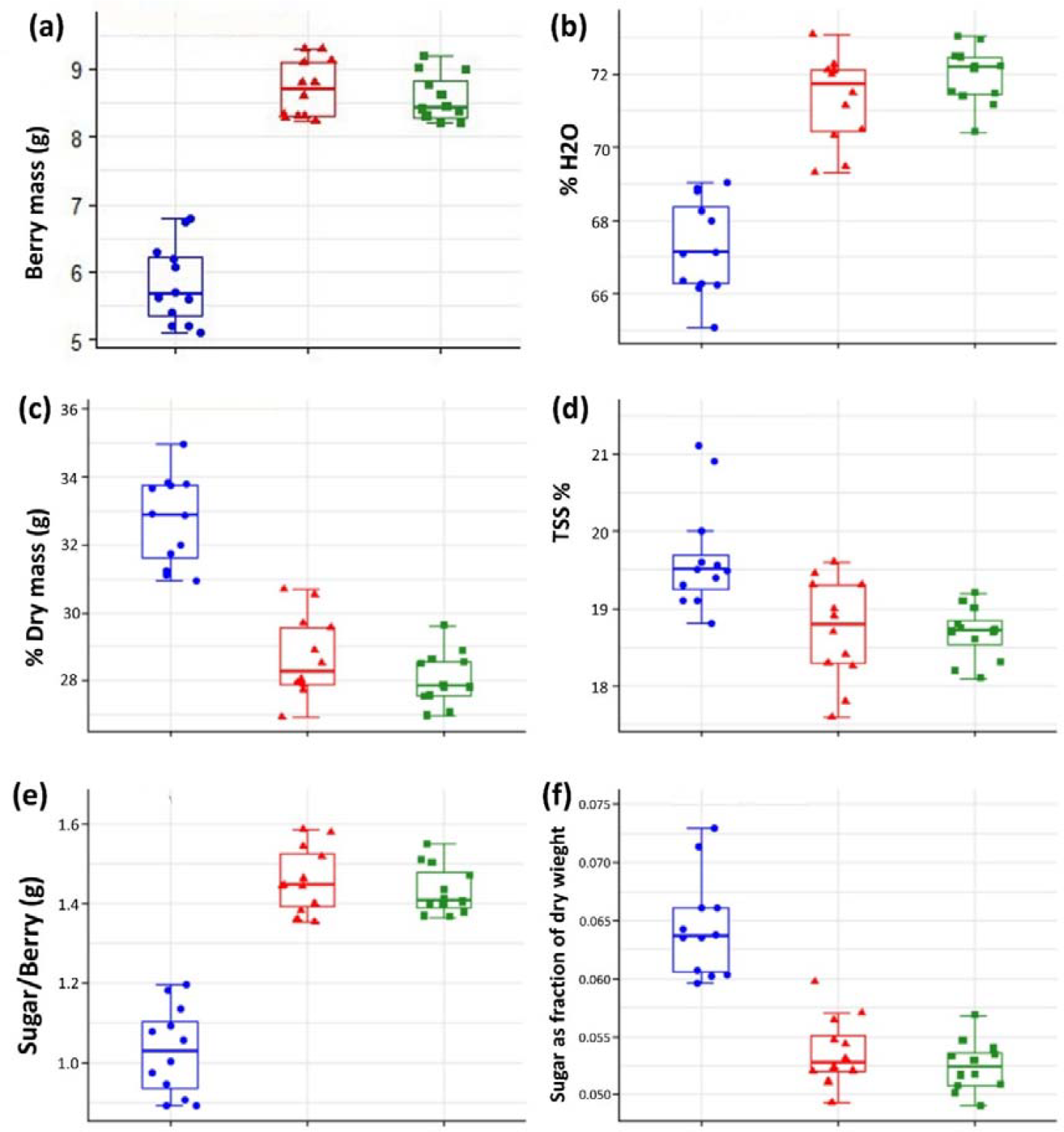
The effect of viral infection on the berry quality among the infected clones (clone 3236 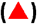 and clone 3215 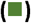) and the control 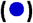, Each clone comprised 6 replicates. **(a)** The effect viral infection on berry fresh weight, (b). The effect viral infection on moisture percentage, **(c)**. The effect viral infection on berry dry weight, **(d)**. The effect viral infection on total soluble solids, **(e)**. The effect viral infection on sugar content of the berry, and **(f)**. The effect viral infection on sugar content as a fraction of the dry weight %., one-way ANOVA has been calculated along with the plot using ggpubr “R” Package.

Depending on the moisture percentage, a significant increase in berry water accumulation can be noticed in the infected clones compared to the control (Table 1, Figure 3(b)). The berry water weight in grams was calculated depending on the moisture percentage. A simple linear regression was calculated to predict the relationship between the berry mass and berry water mass among the control and the infected clones. The linear regression revealed a significant relationship (*F* (1, 34) = 3918, *p* < 0.000001), with a multiple *R*^2^ of 0.9914. It indicates that the berry mass increases significantly in the infected clones depending on the increase in water accumulation in the berry. However, the relationship between the dry components of the berry and the berry mass needs to be clarified. Since the percentage of the dry mass shows a highly significant difference between the infected clones and the control (Table 1 and Figure 3(c)), the dry berry mass is calculated in gram depending on the dry weight percentage. A linear regression was calculated to predict the relationship between berry mass and dry berry mass. A significant relationship was found (*F* (1, 34) = 268.6, *p* < 0.00000), with a significant multiple slope value (*R*^2^ = 0.8867). However, the individual slope for each clone shows that only the control had a high slope value (*R*^2^ = 0.91), while the clones 3236 and 3215 had slope values of 0.6 and 0.66, respectively, indicating that the increase in berry mass was due to ectopic accumulation of water rather than dry mass.

The °*Brix* value can be defined as the mass of sucrose per 100 g of juice, although it measures all soluble solids per 100 g of juice. The MANOVA analysis revealed a significant difference in the °*Brix* value between the infected clones and the control, where no significant effect was noticed of the season on the °*Brix* value (Table 1 and Figure 3(d)). Sucrose typically represents around 95 % of the TSS in grape juice. °*Brix* considers that an acceptable approximate determination of the sugar. Berry sugar was calculated using the method proposed by Vila et al. (2010), (Equation (7)).

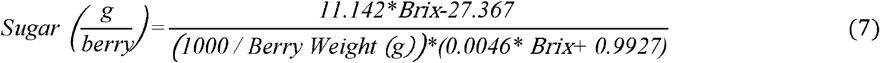

The ANOVA test was conducted to explain the effect of viral infection on the amount of sugar in the berries, showing a statistical significance (F (2,3) = 4.170, P <0.0001). The amount of sugar/ berry was higher in both infected groups relative to the control, although the °*Brix* value was lower in the infected clones than the control, which could be related to the berry mass. Analysis of covariance (ANCOVA) confirmed a significant effect of infection type on the berry sugar content after controlling for berry mass (*F* (3, 32) = 332.9, *P* < 0.001, *R*^2^ =0.969). Figure 4 (a). shows that the relationship is strongly positive and linear. However, previous studies showed that the relationship between the berry sugar content and the berry weight followed the second-order curve (Considine, 2004). Therefore, forcing the line through the origin is reasonable, and as shown in Figure 4 (b), the differences in sugar/berry between the infected and control groups were related to the berry weight.

**Figure 4.**
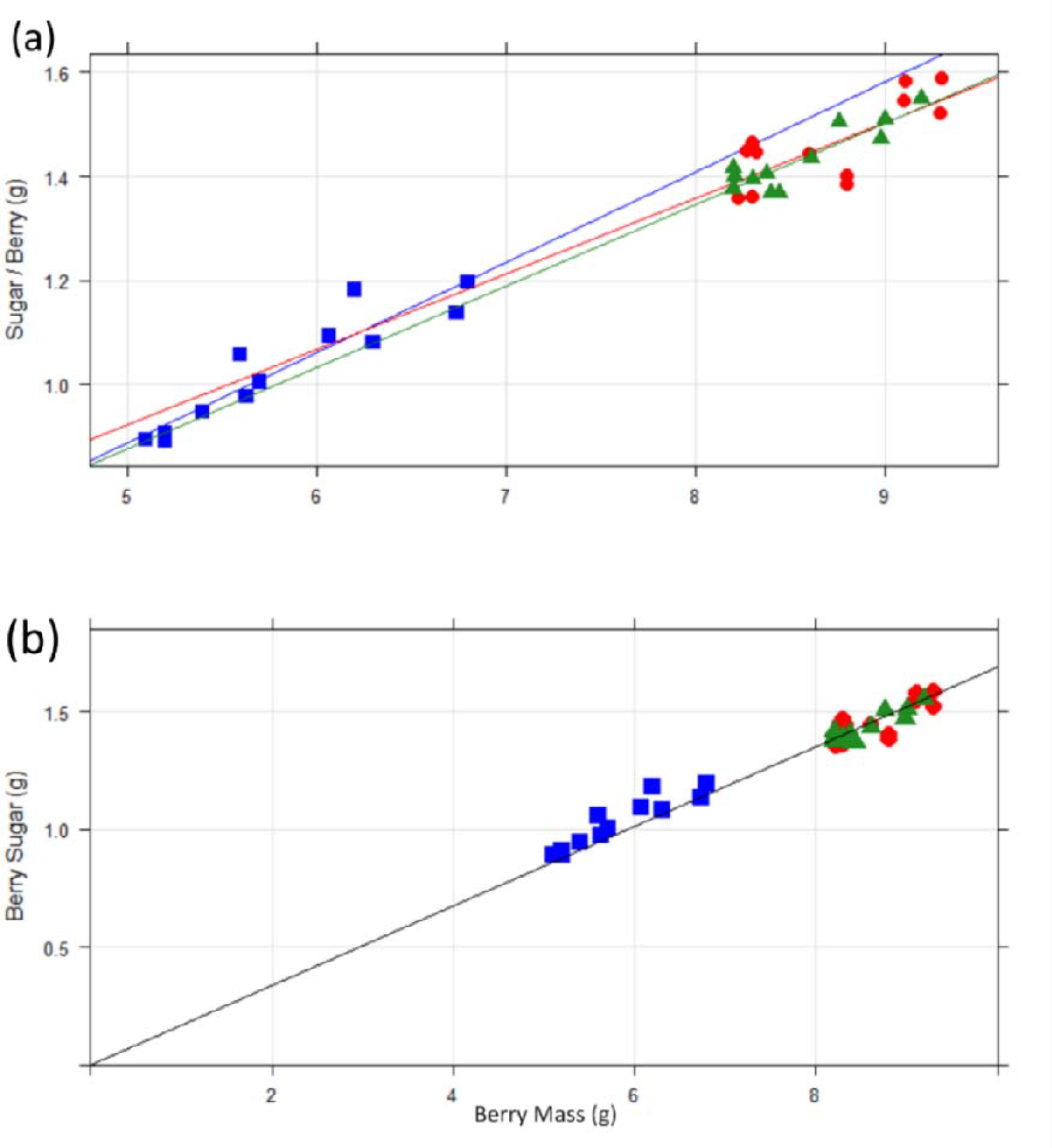
The relationship between the berry sugar mass (g) and average berry mass (g). **(a)**. ANCOVA test of the sugar/ berry (g) tested against the berry mass for each clone, clone 3236 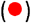, clone 3215 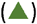 and the control 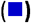 Individual line regression for each infection type. **(b)**. The ANCOVA test show combined relationship by forcing all the lines through the origin point. *F* (1, 35) = 28,989.570, *p* < 0.001, with an *R*^2^ of 0.999, *n*=6 for each clone.

Sugar represents the major component of the TSS. Therefore, analysing the sugar as a fraction of the dry berry mass against the berry mass will clarify the picture of the viral effect on the sugar content. First, the sugar was calculated per berry dry weight as shown in equation (8):

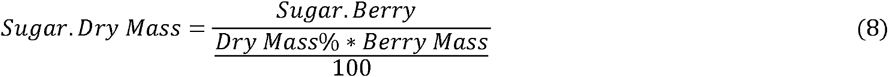

Then the effect of the infection on sugar as a fraction of dry mass was assessed by ANOVA. The results showed a significant difference between the infected clones and the control (*F* (2,32) = 21.4, *p* = 0.001; Figure 3 f)). The value of the sugar as a fraction of the dry mass was higher in the control compared to the infected clones. The result was confirmed by Tukey post-hoc contrasts, which showed a significant difference between control versus clone 3236 (*p* <0.001) and clone 32315 (*p* <0.001), and no significant difference between the infected clones.

Table 1 shows that the TA has a significant difference between the control and the infected clones, whereby TA is lower in the control group than in the infected clones. Moreover, the seasons had a significant effect on the TA (Table 1, Figure 5(a)). Typically, the TSS (°*Brix*) and TA bear a negative relationship throughout the course of ripening. The relationship between TSS and TA provides an indication of fruit maturity and quality (Figure 5 (b)). Interestingly, the two groups of virus infections show a noticeable difference in the regression slope of the TSS: TA (Figure 5(c)). The control shows an inverse relationship between TSS and TA, as expected *y*=37-35x, *R*^2^ =0.55. Clone 3236 reveals almost similar trend to the control, *y*=30-20x, *R*^2^ =0.42; however, the slope was less sharp than the control. In contrast, clone 3215 shows a considerably weaker relationship where the slope was almost flat, *y*=21-3.8x, *R*^2^ =0.032 (Figure 5(c)).

**Figure 5.**
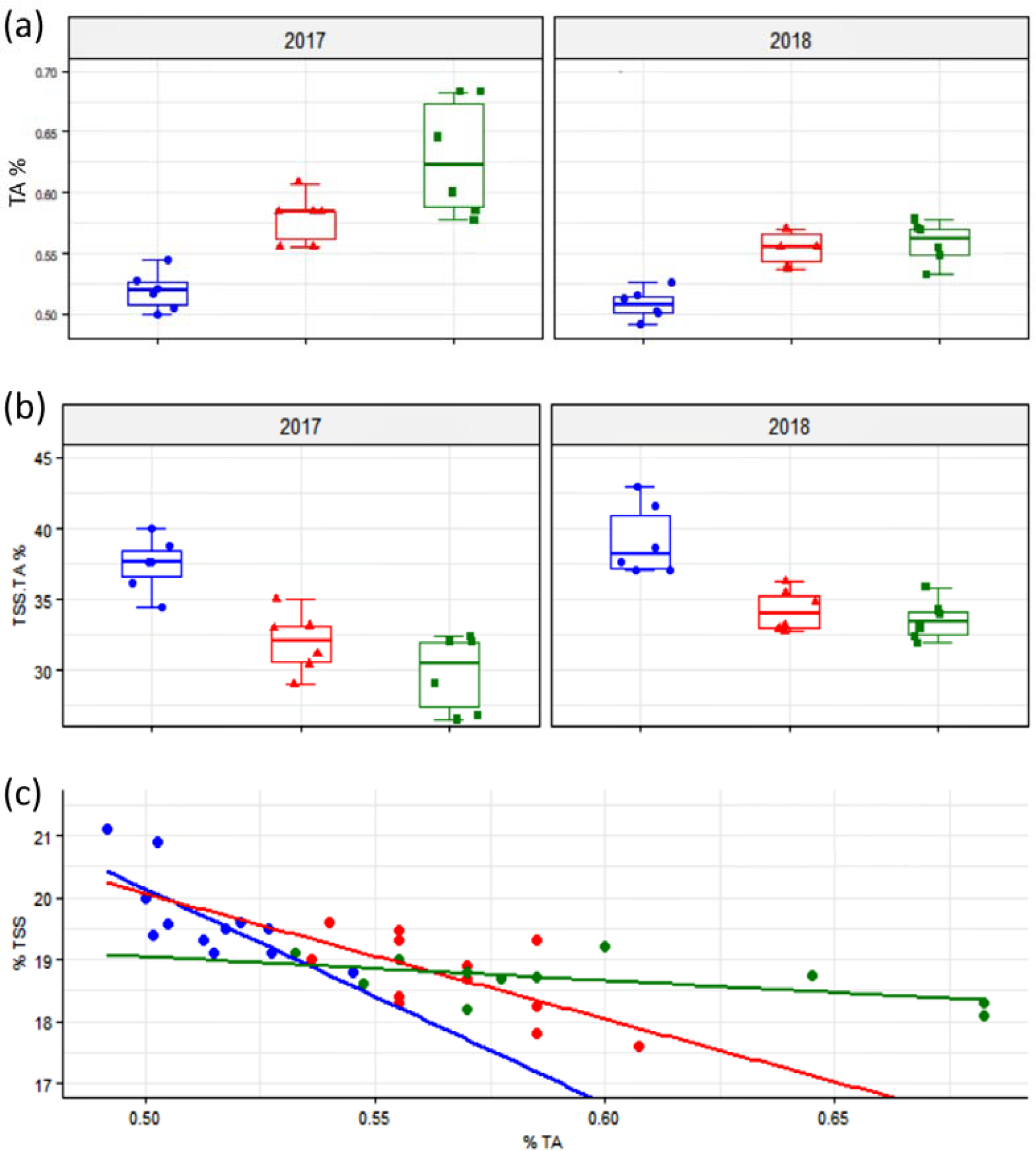
The effect of the viral infection on the acidity (TA) of the grape berry and relationship to TSS between the control 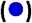 and the infected clones (clone 3236 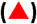and clone 3215 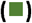). **(a)**. The effect on the TA as a percentage in the berry juice. **(b)**. The effect of viral infection on the TSS:TA ratio. **(c)**. The relationship between the °*Brix* value and the TA among the infection type group.

It is apparent from Figure 2 that virus infection influenced berry colouration. Analysis revealed a significant difference in the amount of anthocyanin among the infection groups (Table 1, Figure 6(a)). Tukey post-hoc contrasts showed that anthocyanin content in clone 3236 and clone 3215 was significantly less than the control (*p* <0.001), but there was no difference between the infected clones. A similar trend was evident in total phenols; however, the magnitude of the difference was less pronounced (Figure 6(b)).

**Figure 6.**
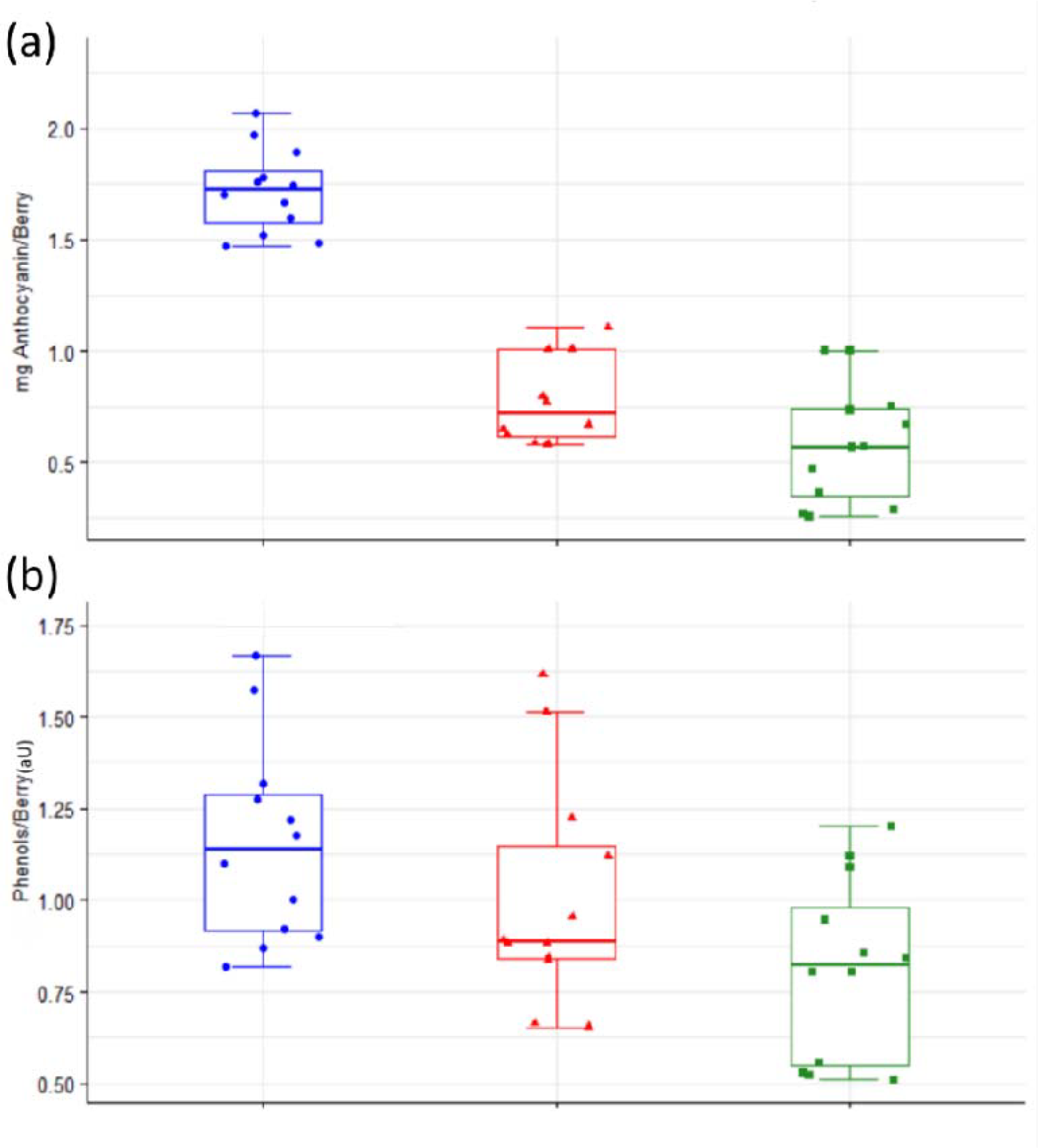
The effect of viral infection on anthocyanin and phenol levels between the control 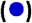 and the infected clones (clone 3236 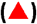 and clone 3215 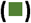). **(a).** The berry anthocyanin content (mg) per berry. **(b).** The phenol content (mg) per berry. AU: absorbance Unit.

### The interaction of viral load with berry quality

Viral load was determined for the four assayed viruses by RT-qPCR. As shown before, the berry mass and anthocyanin were the most affected characteristics by the viral infection. A linear regression of berry mass and the anthocyanin against GLRaV-3 load in both infected clones showed a linear relationship. However, the slope value was low for both seasons, suggesting a weak relationship (Figure 7(a) and (b)). An ANCOVA test was applied using the season as the primary variate and the GLRaV-3 load for both infected clones as a covariate against the berry quality parameters (berry mass, °*Brix*, titratable acidity, dry mass and anthocyanin), to clarify the relationship between the viral load and the berry quality.

**Figure 7.**
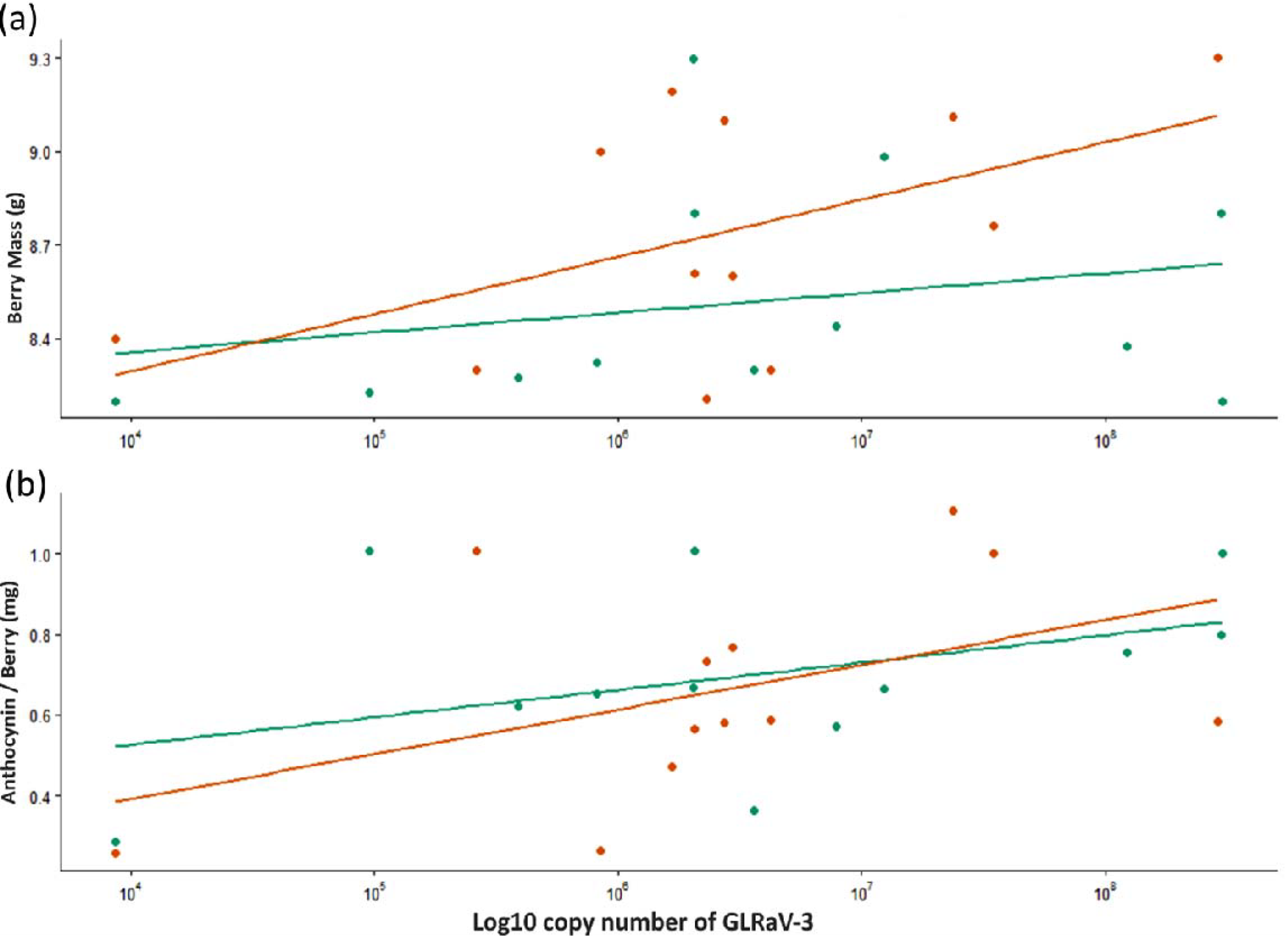
Summary of the linear regression of the viral load. **(a)**. The linear model of the berry’s mass (g) against the log10 of the copy number of GLRaV-3 for two seasons 2017 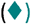, 2018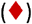. (b). The linear model of the berry’s anthocyanin (mg) against the log10 of the copy number of GLRaV-3 for two seasons 2017 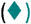, 2018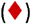, for the infected clones (clone 3236 and clone 3215).

The results showed no significant differences between the GLRaV-3 virus copy number and the measured berry quality parameters. A similar trend was observed for both GLRaV-4 strain 9 and GVA when the ANCOVA test has performed. Interestingly, the results were similar for the three viruses, as clearly indicated by the *p*-value shown in Table 2. Moreover, the results indicated by the *R*^2^ and adjusted *R*^2^, which represent the slope, were very low and mostly null. That refers to the null relationship between the virus copy number and the measured berry quality parameters. On the other hand, the season effect has a considerable impact on titratable acidity, as evidenced by the *p* value of 0.001 for all viruses. Although the *p* values in the three viruses were nearly identical, the slope values (*R*^2^) in GLRaV-3 were slightly lower when compared to GVA and GLRaV-4 strain 9, as shown in Table 2.

**Table 2.** The summary of the ANCOVA test including a season as the primary variate and the GLRaV-3 load for both infected clones as a covariate against the berry quality parameters.

|  | Mass |  | °Brix |  | Titratable acidity |  | Dry Mass |  | Anthocyanin |  |
| --- | --- | --- | --- | --- | --- | --- | --- | --- | --- | --- |
|  | *M* | *p* | *M* | *p* | *M* | *p* | *M* | *p* | *M* | *p* |
| (Intercept) | 8.48 | <0.001 | 18.55 | <0.001 | 0.61 | <0.001 | 28.41 | <0.001 | 0.66 | <0.001 |
| GLRaV-3 | 0 | 0.425 | 0 | 0.949 | 0 | 0.604 | 0 | 0.197 | 0 | 0.314 |
| Seasons [2018] | 0.24 | 0.136 | 0.3 | 0.17 | -0.05 | **0.002\*\*\*** | -0.33 | 0.441 | -0.02 | 0.844 |
| Observations | 24 |  | 24 |  | 24 |  | 24 |  | 24 |  |
| $R^2 / R^2$ adjusted | 0.115 / 0.030 | | 0.091 / 0.005 | | 0.371 / 0.311 | | 0.118 / 0.034 | | 0.054 / -0.036 | |

**GLRaV-4 strain 9**
|  | Mass |  | °Brix |  | Titratable acidity |  | Dry Mass |  | Anthocyanin |  |
| --- | --- | --- | --- | --- | --- | --- | --- | --- | --- | --- |
|  | *M* | *p* | *M* | *p* | *M* | *p* | *M* | *p* | *M* | *p* |
| (Intercept) | 8.33 | <0.001 | 18.78 | <0.001 | 0.62 | <0.001 | 28.64 | <0.001 | 0.73 | <0.001 |
| GLRaV-4 | 0 | 0.375 | 0 | 0.132 | 0 | 0.263 | 0 | 0.067 | 0 | 0.124 |
| Season [2018] | 0.27 | 0.197 | 0.12 | 0.535 | -0.07 | **0.007\*\*\*** | -0.32 | 0.436 | -0.05 | 0.757 |
| Observations | 12 |  | 12 |  | 12 |  | 12 |  | 12 |  |
| $R^2 / R^2$ adjusted | 0.247 / 0.079 | | 0.254 / 0.088 | | 0.597 / 0.508 | | 0.365 / 0.224 | | 0.253 / 0.086 | |

**GVA**
|  | Mass |  | °Brix |  | Titratable acidity |  | Dry Mass |  | Anthocyanin |  |
| --- | --- | --- | --- | --- | --- | --- | --- | --- | --- | --- |
|  | *M* | *p* | *M* | *p* | *M* | *p* | *M* | *p* | *M* | *p* |
| (Intercept) | 8.46 | <0.001 | 18.43 | <0.001 | 0.64 | <0.001 | 27.72 | <0.001 | 0.45 | 0.005 |
| GVA | 0 | 0.709 | 0 | 0.071 | 0 | 0.355 | 0 | 0.03 | 0 | 0.062 |
| Season [2018] | 0.24 | 0.327 | 0.3 | 0.167 | -0.08 | **0.007\*\*\*** | 0.15 | 0.731 | 0.1 | 0.54 |
| Observations | 12 |  | 12 |  | 12 |  | 12 |  | 12 |  |
| $R^2 / R^2$ adjusted | 0.187 / 0.007 | | 0.336 / 0.189 | | 0.578 / 0.485 | | 0.460 / 0.340 | | 0.344 / 0.198 | |

\*\*\**p* < 0.001, \*\**p* < 0.01, \**p* < 0.05, *p* < 0.1, M: Mean, *p*: *p* value

## 4 Discussion

Berry quality is an essential standard in table grape production since it is served fresh to the consumers (Rolle et al. 2012). Berry ripening progresses in two growth stages consisting of berry formation and maturation, separated by a lag phase coinciding with véraison (Coombe and McCarthy, 2000). Previous studies indicated that the pre-véraison berries from virus-infected vines do not show a significant difference in quality compared to virus-free vines (Alabi et al. 2016). In contrast, dramatic differences were observed during post-véraison, suggesting that viral infection caused more significant impacts on ripening-related processes starting from véraison (Alabi et al. 2016; Montero et al. 2016; Montero et al. 2016a). As a result, this study concentrated on the final stage of growth, when the virus full effect could be clearly observed on the berries.

Berry size is an essential feature in both wine and table grapes. In wine grapes, smaller berries are desired, as the anthocyanin and sugar will be more concentrated in the small berry volume (Abu-Zahra, 2010; Chen et al. 2018a; Ferrer et al. 2014; Melo et al. 2015; Nuzzo and Matthews, 2005; Weaver and Winkler, 1952). However, large berries are more desirable in table grapes considering the berry’s density (Río Segade et al. 2013). Increased berry size is the most obvious indication of viral infection in the cv. Crimson Seedless vines (Singh Brar et al. 2008). The berry’s size, or fresh weight, is regulated by cell number, cell volume, and the accumulation of organic substances (sugars) in the cell vacuoles (Coombe, 1976; Ollat et al. 2002; Robinson and Davies, 2000). Thus, the pulp cells enlarge as a result of an influx of sugars and water into the vacuole. The vacuoles of the pulp cells form about 99% of the cell volume (Diakou and Carde, 2001). In the virus-free vines, pulp cells continued to enlarge throughout ripening, undergoing significant structural changes responsible for berry softening late in the harvesting stage (Barnavon et al. 2000; Nunan et al. 2001). In comparison, virus-infected vines frequently exhibit a delay in ripening, one of the most obnoxious symptoms of viral infection (Martínez et al. 2016; Over de Linden and Chamberlain, 1970). A delay in ripening is associated with a prolonged influx of water and sugar into the berry from the leaves. The water and sugar will accumulate in the mesocarp cell vacuoles, leading to an increase in berry weight (Bobeica et al. 2015; Fontes et al. 2011; Rowhani et al. 2015). The increase in berry size has been clearly observed in the present study (Figure3(a)).

At the same time, the ripening of grape berries is accompanied by sugar accumulation; these processes play significant roles in the quality of the berries. Sugars are accumulated in the mesocarp vacuoles, which account for 65–91% of the fresh weight of a ripened berry (Fontes et al. 2011; Marty, 1999; Pastore et al. 2011). However, the results show a reduction in the TSS that was measured as °*Brix* value in both infected vines; GLRaV-3 implied a remarkable decrease in TSS. It could be argued that the fact of sugar accumulation in the berry is controlled by a feedback inhibition mechanism, whereby sugar controls the expression of sugar transporter genes (Koch, 1996; Lecourieux et al. 2014). The feedback mechanism keeps the sugar level balanced with the berry mass (Lemoine et al. 2013). As evident in Figure 4(b), sugar was the major contributor to the increase in berry weight. However, the °*Brix* value was low in the infected berry as a result of the dilution of the sugar concentration by the increase in weight due to water.

Tartaric and malic acids are the main organic acids in the berry (Cholet et al. 2016; Kliewer et al. 1967; Lamikanra et al. 1995). Both acids increased rapidly during the pre-véraison stage, reaching their highest levels near véraison, and then declined throughout ripening (Muñoz-Robredo et al. 2011). However, the data shows that the TA% was significantly higher in both infected clones, most notably in the mixed clone Figure 5(a), which is a common sign of GLRaV-3 infection (Alabi et al. 2016; Kliewer and Lider, 1976). The high level of TA in the infected vines was associated with delayed ripening (Martínez et al. 2016). The decline of organic acids is controlled by many factors, such as enzymatic degradation (Batista-Silva et al. 2018; Lakso and Kliewer, 1975; Sweetman et al. 2014) the dilution effect, the increase in temperature and the acid salt formation (Kliewer et al. 1967; Ruffner, 1982). Among these, the acid salt formation might explain the delay in the decline of the organic acids in grape berries. Inorganic salts are transported from the root to the leaves through the xylem at the first stage of berry development. After véraison, the salts were transported from the leaves and unloaded into the berries along with sugar via the phloem. The salt content of the berry will decrease because the virus disrupted phloem transport (Ford, 2012; Jayasena and Cameron, 2008; Kliewer, 1966). The relatively low salt content of virus-infected berries will prevent acid salt formation and keep the acidity higher in the infected berries. Interestingly, the seasonal effect on both TA and pH was significant. Similar results were previously reported (Lee et al. 2009; Lee and Martin, 2009), which may have resulted from the influence of different temperatures over the two years; however, this was not explored here (Lakso and Kliewer, 1975).

Previously, two types of phenolic chemicals found in berries were investigated: tannin and anthocyanin (Adams, 2006). At the initial stage of berry formation, tannin chemicals were collected in the mesocarp. By contrast, anthocyanin accumulates in the skin following véraison (Kennedy et al. 2007). Evidently, anthocyanin is more susceptible to virus infection than other forms of phenols. The result has demonstrated a significant decrease in anthocyanin concentration in infected clones compared to control vines, which was particularly pronounced in clone 3215 (mixed infected clones) (Figure 6(a)). Additionally, the reduction was extended to phenolic compounds and demonstrated the same mechanism, but the difference was less pronounced between the control and the infected clones (Figure 6(b)). Although sugars accumulation in skin is very low compared to the pulp, the evidence is growing for some control of polyphenol metabolism by sugars, possibly through crosstalk with genes responsible for sugar regulation (Dai et al. 2013; Smeekens et al. Matsushima et al. 1989; 2010; Solfanelli et al. 2006; Zheng et al. 2009). Previous studies show that sugars modulate anthocyanin biosynthesis pathway genes (Filippetti et al. 2015; Lecourieux et al. 2014; Solfanelli et al. 2006). Beyond the effect of delayed ripening, the viral infection seems to have a deleterious influence on anthocyanin synthesis, as indicated by the previously reported up-and down-regulation of several particular genes (Vega et al. 2011).

Moreover, research indicates that phloem-specific viruses disturb the leaf minor vein phloem transporting process (Naidu et al. 2015). Disturbing phloem vines leads to aggravating sugar transport from the source (leaves) to the sink (berries) (Pawar and Rana, 2019), resulting in an accumulation of sugar in the leaf mesophyll cells, which influences anthocyanin production, leading to red colouration of interveinal areas of infected leaves. These findings are consistent with studies indicating increased levels of anthocyanins in symptomatic leaves of GLD-affected red-berried grapevines (Gutha et al. 2010). Although aspects of the host-virus interaction between GLRaV’s and grapevine remain ambiguous, it appears the major influence of the virus on berry quality results from an altered source/sink balance, which in particular affects sugar transport in the phloem vessels after véraison (Alabi et al. 2016; Naidu et al. 2015). The data presented here is consistent with this theory. Moreover, it provides no evidence that viral load is important, but rather the virus proteins play an essential role in reprogramming gene expression. Additionally, the findings reveal that clone 3215 exhibits a modest increase in berry quality variation compared to clone 3236. This might be explained as the combined effect of mixed viral infections in clone 3215. However, no statistically significant evidence was found between both clone infections, and most of the effect would belong to the case where GLRaV3 is present.

## Notes

### Competing Interest Statement

The authors have declared no competing interest.

